# Valproic Acid Affects Neuronal Specification and Differentiation During Early Optic Tectum Development of Zebrafish

**DOI:** 10.1101/2022.06.15.496299

**Authors:** Sierra C. Dixon, Bailey J. Calder, Shane M. Lilya, Brandon M. Davies, Annalie Martin, Maggie Peterson, Jason M. Hansen, Arminda Suli

## Abstract

The mammalian superior colliculus and its non-mammalian homolog, the optic tectum (OT), are midbrain structures that integrate multimodal sensory inputs and guide non-voluntary movements in response to prevalent stimuli. Recent studies have implicated this structure as a possible site affected in Autism Spectrum Disorder (ASD). Interestingly, fetal exposure to valproic acid (VPA) has also been associated with an increased risk of ASD in humans and animal models. Therefore, we took the approach of determining the effects of VPA treatment on zebrafish OT development as a first step in identifying the mechanisms that allow its formation. We describe the normal OT development during the first 5 days of development and show that in VPA treated embryos, while proliferation of the OT neuroepithelium continued, neuronal specification stalled. This was followed by impairment of neurite extension and complexity, suggesting that in addition to neurogenesis, VPA treatment affects axonogenesis and dendritogenesis. VPA treatment was most detrimental during the first three days of development and did not appear to be linked to oxidative stress. In conclusion, our work provides a foundation for research into mechanisms driving OT development, as well as the relationship between the OT, VPA, and ASD.

## INTRODUCTION

The superior colliculus (SC) is a mammalian midbrain structure that receives visual, auditory, and somatosensory inputs. It is known for its ability to direct movement of the eyes ^1^, ears ^2^, and limbs ^3^ towards salient events in the environment. Engagement of the SC in cross-modal sensory integration leads to an increased ability to detect events ^4,5^, a faster reaction time ^6-11^, and an increased ability to localize targets or events ^11,12^. Ablation studies have shown that while the cortex allows for discrimination of sensory inputs, the SC is necessary to initiate movements toward these events ^13^. Recently, this subcortical area has been implicated in influencing social behaviors and has been purported to lead to neurodevelopmental disorders such as Autism Spectrum Disorder (ASD) ^14-17^. Although many studies have shed light on SC function, its microcircuitry and formation continue to be poorly understood.

Work utilizing the genetically-tractable zebrafish model organism has become instrumental in dissecting the microcircuitry of the optic tectum (OT), the non-mammalian homolog of the SC. Similar to the SC, the OT receives sensory inputs and is involved in phototaxis, hunting, and predator avoidance ^18-24^. During embryonic and larval development, the OT consists of the periventricular layer (PVL), where the neuronal cell bodies reside, and the neuropil, where the neurites extend ^18^. To better understand OT development, we exposed embryos to a known developmental neurotoxin, valproic acid (VPA), with the intention of identifying the processes, and eventually the molecular underpinnings, required for proper OT development. Valproic acid was chosen due to its use as a model for ASD in zebrafish as characterized by deficits in both social communication and interactions, as well as restricted and repetitive behaviors, interests, and activities. Both genetic and environmental factors have been linked to ASD etiology, and treatment of zebrafish embryos with VPA has been shown to induce similar social interaction deficits ^25-28^ and affect ASD-associated genes ^27^.

Valproic acid is a commonly used drug in the treatment of epilepsy, bipolar disorder, and schizophrenia; however, in 1984 it was shown to have adverse effects in the developing fetuses of pregnant mothers taking VPA. These effects were termed “fetal valproate syndrome” ^29^ with ASD being one of the most prominent disorders associated with it. Results from early, smaller, studies showed that VPA monotherapies correlated with much higher rates of ASD (12% rate) compared to unexposed groups (1.9% rate) and were significantly higher than other antiepileptic drugs (both carbamazepine and lamotrigine), which resembled unexposed groups ^30^. As more information becomes available, including that from larger studies, data shows that VPA exposure not only affects morphological endpoints during development, such as neural tube defects, but is also associated with the prevalence of ASD, confirming the results of the earlier studies. In fact, the risk of developing ASD for individuals born to mothers exposed to VPA during pregnancy is fivefold higher than the general population ^31^. Mechanisms of VPA-induced developmental toxicity are not fully understood, but potential contributors include VPA-induced oxidative stress and histone deacetylase (HDAC) inhibition.

In our study, we exposed zebrafish embryos to VPA from the onset of gastrulation, 6 hours post fertilization (hpf), to 5 days post fertilization (dpf) and focused specifically on OT development. We show that VPA exposure leads to stalled neurogenesis and perturbation of neurite development. Additionally, we identify a critical window for the effect of VPA exposure on OT. Finally, we demonstrate that these effects do not seem to be caused by oxidative stress, since pretreatment with a drug that induces protective stress-response genes does not ameliorate the effects of VPA.

## MATERIALS AND METHODS

### Zebrafish Lines and Husbandry

All experiments were performed according to IACUC approved protocols at Brigham Young University (IACUC Protocol Number: 19-0901). Embryos and larvae were raised on a 14h:10h light:dark cycle.

The *y304Et(cfos:Gal4); Tg(UAS:Kaede)* enhancer trap line ^32^ was generously provided by the lab of Harold Burgess at NIH.

The *Tg(NeuroD:tRFP*^*w68*^) line was generated using the Multisite Gateway system ^33,34^ to make the NeuroD:tRFP construct from 1) pME NeuroD (a gift from Teresa Nicolson containing the 5kb region of *neurod1* 5’promoter ^35^ 2) pME tagRFP (a gift from Chi-Bin Chien) and 3) p3E polyA (Multisite Gateway system). The construct was injected at one-cell stage embryos and stable transgenic were subsequently isolated.

### Embryo Medium (EM)

EM was made from 20XE2 (17.5g NaCl, 0.75g KCl, 4.9g MgSO4-7H2O, 0.41 g KH2PO4, 0.12g NaH2PO4-2H2O, into water for 1L of total solution).

### Chemical Treatments

A 250mM VPA stock solution was made by dissolving sodium valproate salt (Sigma, P4543) into EM. A working solution of 250µM VPA in EM was prepared from the stock solution. The EM was carefully maintained at pH 7.20. A stock solution of 12.5mM D3T(Sigma, D5571) in DMSO was made. A working solution of 5µM D3T was then made by dissolving the D3T stock solution in EM. The final concentration of DMSO in the working solution was 0.04%.

A new VPA stock and working solution was made and administered daily at 5pm beginning at 6hpf. 10 embryos/well were incubated in 6-well plates; three wells received VPA treatments and three received only EM. Each well contained 7ml of solution. Every day a new solution was made, and the old solution was replaced. The single dose of D3T was administered from 6hpf to 18hpf concurrently with 250µM VPA, or as a pretreatment without VPA, then washed out and replaced with a 250µM VPA only solution. Embryos were then treated daily with VPA as previously outlined.

### Confocal Microscopy

Embryos and larvae were screened for expression of the fluorescent reporter, and 3 embryos/condition were embedded dorsal-side down in 1.5% low melting point agarose (LMA) (Gene Mate, E-3126-125) with 1:25 MESAB (Syndel, 200-226) in 10mm glass-bottom microwells. Images were obtained on an Olympus Fluoview FV1000 confocal microscope using a 488nm laser (*y304Et(cfos:Gal4); Tg(UAS:Kaede))* or a 546nm laser *(Tg(NeuroD:tRFP*^*w68*^)) with a 20X lens. Images were collected at 640 × 640 pixels at 2μs/pixel. The HV of the 543nm laser was held constant across wild-type and treated embryos, so fluorescence could be compared between the two.

For time-lapses, multiple embryos (between n=3 and n=5) were anesthetized and embedded dorsal-side down in 14mm glass-bottom microwells. Valproic acid treated embryos were embedded in 1.5% LMA,1:25 MESAB, and 250µM VPA. Embedded control and VPA treated embryos were covered with 3mL of the respective solution (EM or 250µM VPA solution). Each embryo was imaged once every ten minutes.

### Photoconversions

Following collection, embryos were raised in the dark to prevent off-target photoconversion of Kaede. After larvae were prepared for imaging as previously described, photoconversion of Kaede was performed utilizing a 405nm laser at 1% power for 5 seconds with a 40X objective lens. Following dispersion of photoconverted Kaede, images were taken with a 40X objective lens and a digital zoom of 6X (3dpf) or 3X (4-5dpf). All other settings for image capture remained the same as previously described.

### Enucleations

At 30hpf embryos were dechorionated, anesthetized and embedded ventral-side down in 1.5% LMA. Using a sharpened tungsten needle, an incision was made in the epithelium anterior of the optic cup, and the optic cup and lens were gently pushed out. The embryos were un-embedded, returned to a dish with fresh EM, and grown to 5dpf in standard conditions.

### Glutathione/glutathione disulfide redox potential analysis

At 6hpf embryos were treated with DMSO (control vehicle) or D3T (5µM). At 18hpf, D3T was removed and VPA (250 µM) was added to half of the wells. At the indicated time points, 30 embryos were collected in 325µL 5% perchloric acid and 0.2M boric acid containing the internal standard γ-glutamylglutamate (10µM) and stored at -20°C. Glutathione and glutathione disulfide were measured with high performance liquid chromatography (HPLC) with fluorescence detection using γ-glutamylglutamate as an internal standard for each sample. Samples were derivatized to S-carboxymethyl, N-dansyl derivatives using the method described^36^. GSH E_h_, GSH and GSSG concentrations were used in the Nernst equation E_h_ = E_o_ + (RT/nF)ln([GSSG]/[GSH]^2^).

### RT-qPCR

At 6hpf embryos were treated with DMSO (control vehicle) or D3T (5µM). At 18hpf, 30 embryos were collected in 250uL TRIzol reagent (Invitrogen) and stored at -20°C. RNA was extracted using a Direct-zol RNA MiniPrep Kit (Zymo Research). cDNA was synthesized via reverse transcription using an iScript cDNA Synthesis Kit (Bio-Rad Laboratories) and used for quantitative real-time PCR (RT-qPCR) analysis using SYBR Green Detection Master Mixes (SABiosciences) on a StepOnePlus real-time PCR cycler (Applied Biosystems). All steps were performed per manufacturer’s instructions. Specific forward and reverse primers for *Gclc, Hmox1a, Hmox1b, Nqo1, Gstp1, and β-actin* were synthesized by Integrated DNA Technologies (Table 1). *β-actin* was used as a housekeeping gene for all samples for normalization purposes. Relative levels of mRNA expression were calculated using the ΔΔC_T_ method relative to the expression of *β-actin* mRNA expression.

**Table 1.**
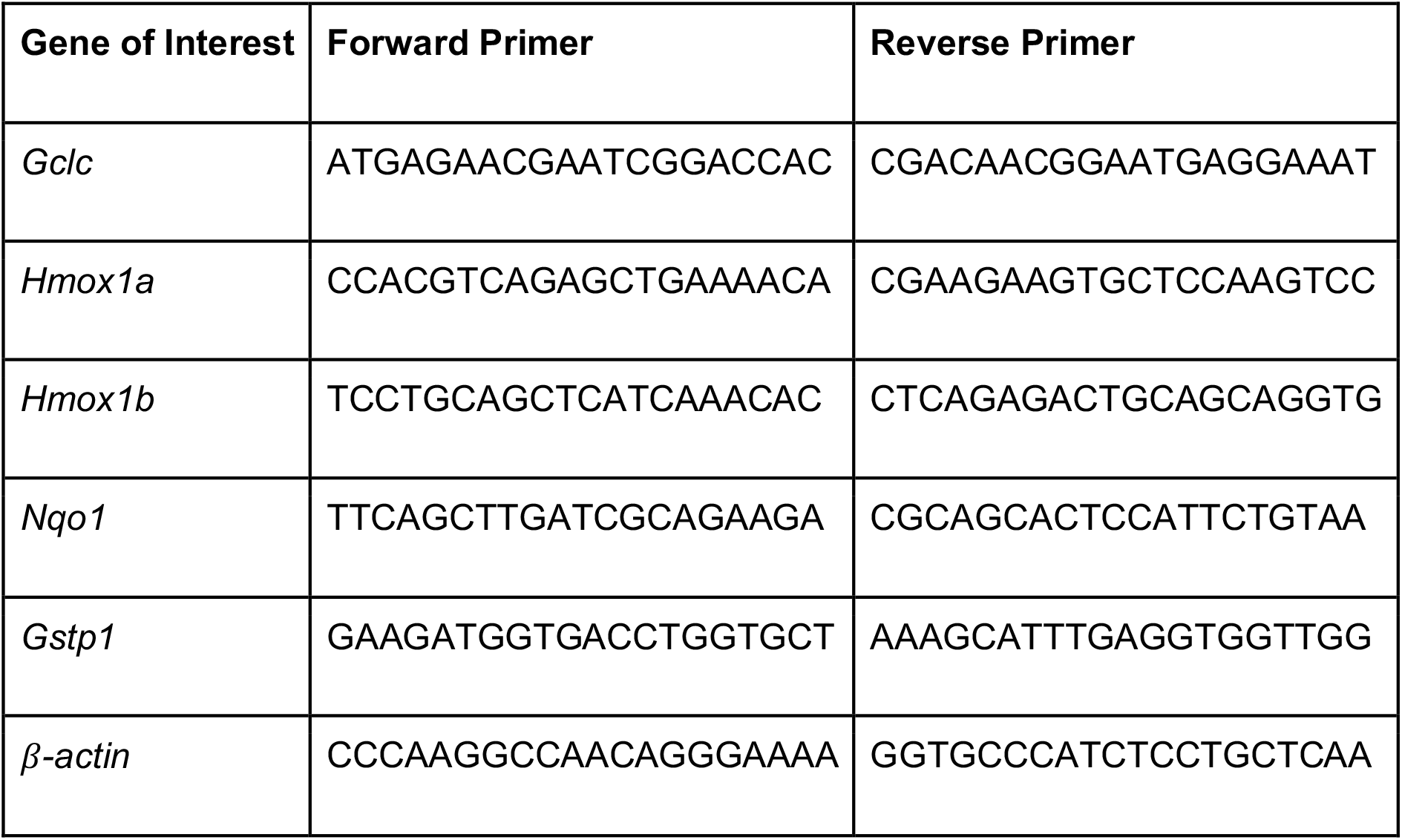
Primer sequences used for amplification of target genes downstream of the nrf2 antioxidant response pathway.

## RESULTS

### Morphological Changes in the OT Following VPA Exposure Reveal Developmental Delays

In order to determine the optimal VPA dose, which would perturb OT development without inducing high mortality rates, dose-response experiments were performed at various concentrations ranging from 0-2000µM VPA (Fig 1A,B). Embryos were treated continuously from 6-120hpf, and the VPA solution was replaced every 24 hours. Embryo survival and hatching was recorded daily, with 250µM identified as the maximum tolerated dose (MTD) due to its high survival and hatching rates. *Note: For our purposes zebrafish from 0hpf-71hpf will be termed embryos and from 72hpf-120hpf will be termed larvae. For any reference that spans the entirety of 0hpf-120hpf we will use the term embryo*

**Figure 1.**
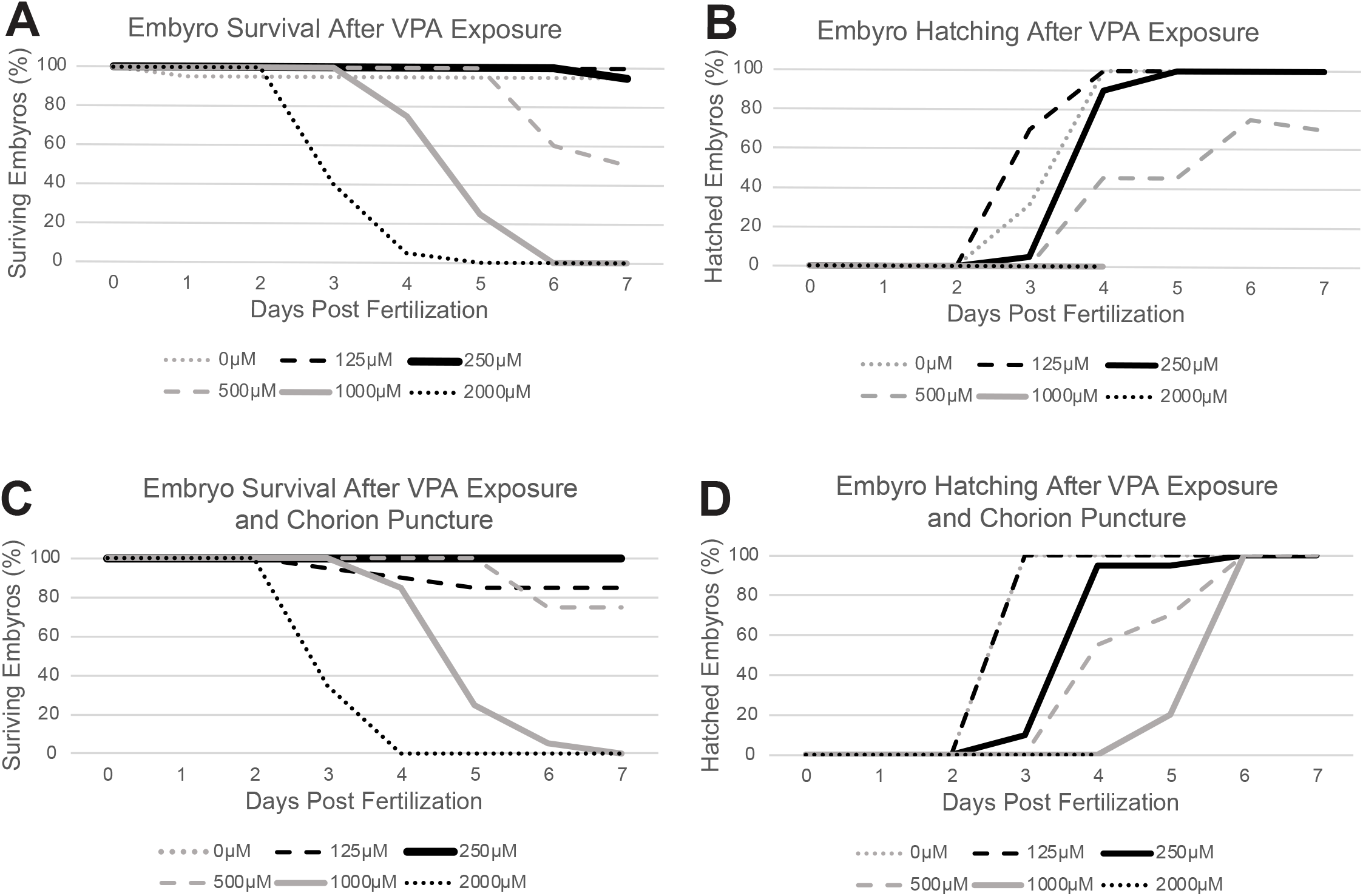
Dose-response experiments identify 250μM VPA as the maximum tolerated dose (MTD). Embryo survival (A) and hatching (B) were recorded daily from 0-7dpf using concentrations of VPA ranging from 0-2000μM. Treated embryos were continuously exposed to VPA from 6-120hpf. 250μM was found to be the MTD due to high embryo survival (A) and hatching (B). (C-D) Experiments repeated in embryos with a chorion puncture showed no appreciable difference in survival or hatching at 250μM.

The *y304Et(cfos:Gal4); Tg(UAS:Kaede)* enhancer trap line ^32^ labels the OT, as well as epiphysis, habenulae, heart, and the olfactory bulbs (sparsely). Therefore, this transgenic line was utilized to determine the morphological effects of VPA treatment during OT development. *Y304Et(cfos:Gal4); Tg(UAS:Kaede)* embryos were treated continuously with 250µM VPA from 6-120hpf (Fig 2A). Valproic acid was replaced every evening and embryos were imaged the following morning from 24-120hpf.

**Figure 2.**
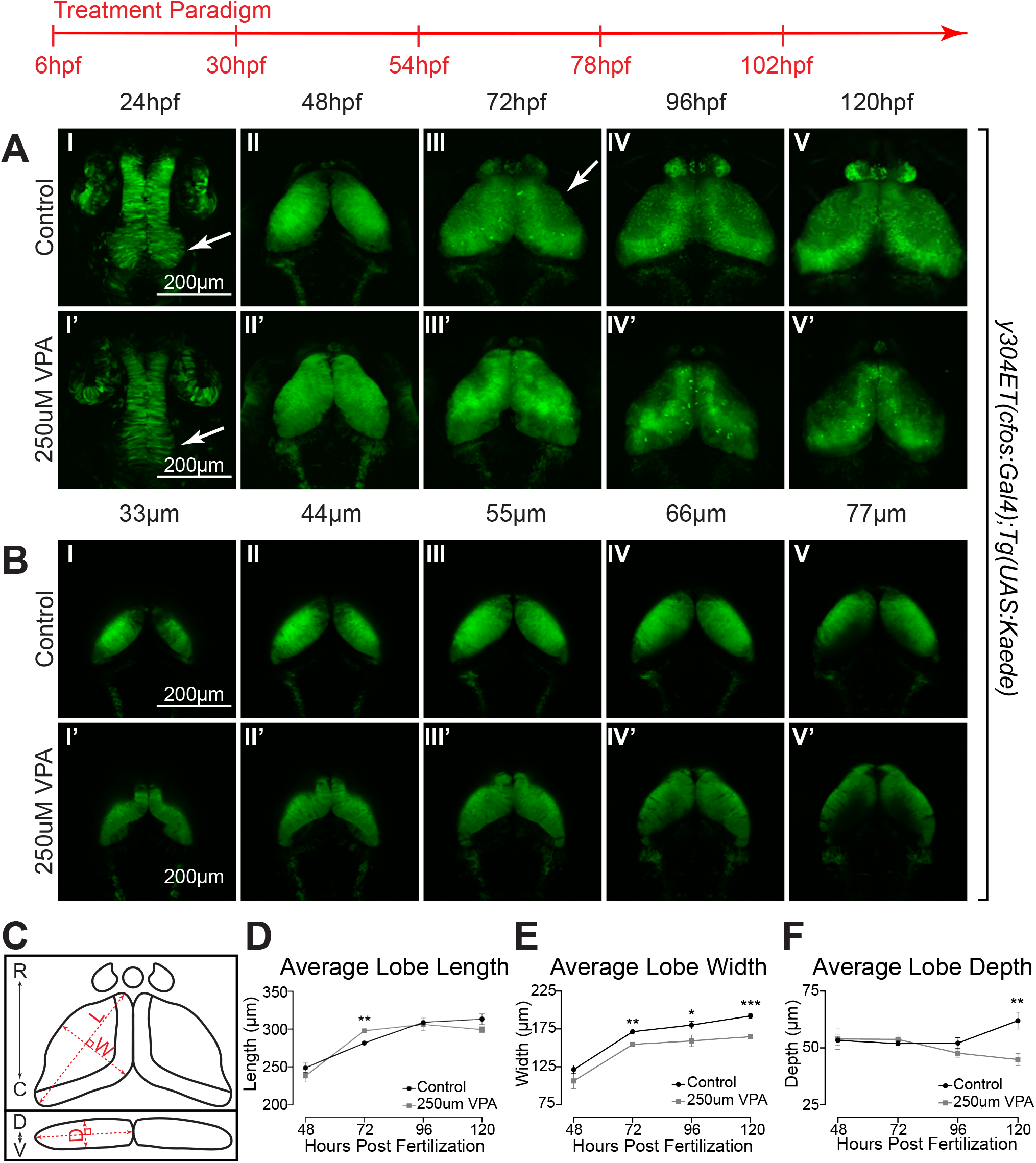
Continuous Valproic Acid treatment causes delayed OT development and decreased neuropil. (A) Daily images of control (I-V) and 250µM VPA-treated *y304Et(cfos:Gal4); Tg(UAS:Kaede)* embryos (I’-V’). Treated embryos were continuously exposed to VPA from 6-120hpf. At 24hpf, proliferation in the neuroepithelium giving rise to the OT is seen in control (arrow, I) but not in treated embryos (arrow, I’). Beginning at 72hpf, the OT neuropil, where the neurites extend, becomes noticeable in control (arrow, III) but not in treated larvae (III’). (B) Images of single slices of dorsal to ventral positions at 48hpf in OT of control (I-V) and VPA-treated embryos (I’-V’). Neurogenesis in treated embryos is delayed as shown by the presence of columnar neuroepithelial cells dorsally in treated embryos (I’-III’) but not in control embryos where neuroprogenitors and early-born neurons (rounded cells) have taken their place (I-III). (C) Cartoon of OT lobe measurements of length, width, and depth. (C) The lobe length of treated larvae is significantly longer than controls at 72hpf but is resolved by 96hpf. (D) Lobe width of treated embryos is significantly increased at all time points, while depth is significantly increased at 120hpf. (Student’s t-test).

At 24hpf, control embryos exhibit a thickening of the neuroepithelium at the posterior part of the neural tube, indicating cell proliferation and the onset of neurogenesis, which is absent in VPA-treated embryos (Fig 2A I,I’, arrows). By 48hpf, z-projected images of both control and VPA-treated embryos display similar OT formation (Fig 2A II,II’). However, individual images show that dorsally the OT of VPA-treated embryos exhibit persistent neuroepithelium (columnar cells) undergoing neurogenesis (Fig 2B I” -III’), while ventrally neuroprogenitor cells and newly born neurons (rounded cells) are present (Fig 2B IV’,V’). In contrast, control embryos do not show the presence of neuroepithelium at any dorsal to ventral position, indicating that neurogenesis is well underway (Fig 2B I-V). The developmental delay became even more apparent at 72hpf when the control larvae show the beginning of neuropil formation (Fig 2A III arrow), indicating that neurons are undergoing axonogenesis and dendritogenesis. The neuropil in VPA-treated larvae seems to be absent at 72hpf and is smaller both at 96hpf and 120hpf. For a more quantitative assessment of the phenotype, we measured the length, width, and depth of each OT lobe (Fig 2C). Valproic acid-treated larvae showed decreased lobe width from 72hpf-120hpf, further evidencing a decrease in neuropil volume (Fig 2E), and decreased lobe depth (Fig 2E). These observations suggest that decreased neurogenesis and prevention of neuropil formation may be two mechanisms by which VPA alters OT development.

In addition to concentration, previous findings have shown the importance of pH in modulating the effects of VPA and its uptake into the cell ^37^. In agreement with these studies, we found that the same dose of VPA at lower pH (pH=6.6) exhibited stronger morphological phenotypes than those at a neutral pH (pH=7.2). In comparison, those at higher pH (pH=7.8) displayed weaker phenotypes (Suppl. Fig 1). To keep the phenotype consistent all subsequent experiments were carried out at a neutral pH (pH=7.2) which was chosen because of its proximity to biological pH and because it is neutral enough to not alter the effects of VPA treatment.

### VPA Delays Neurogenesis in the Developing OT

To better understand the effect of VPA exposure on neurogenesis we utilized the *Tg(NeuroD:tRFP*^*w68*^) transgenic line which drives tRFP via the *NeuroD1* enhancer. *NeuroD1* has been shown to be involved in neuronal differentiation, specification, and migration within the developing mouse cortex ^38^, making it a good indicator of neurogenesis. Additionally, single-cell RNA sequencing (scRNA-seq) data previously collected in our lab ^39^ confirmed the presence of *NeuroD1* in the zebrafish OT and showed that its expression is sequestered to specific populations by 7dpf (Suppl. Fig 2). To capture initial alterations in neurogenesis before large divergences in morphology occurred, we ran two consecutive time-lapses which in their entirety covered 22.5-43.5hpf (Fig 3). These timepoints were chosen because, as mentioned above, between 24-48hpf the neuroepithelium, from which the OT is derived, undergoes initial neurogenesis and gives rise to newly born neurons (Fig 2A I,II,I’,II’, 2B I-V, I’-V”). Timelapses were run on both *y304Et(cfos:Gal4); Tg(UAS:Kaede)* (Suppl. Videos 1-4) and *Tg(NeuroD:tRFP*^*w68*^) embryos (Suppl. Videos 5-10) following the same dosing paradigm used previously to allow for visualization of cell movement and morphology as well as identify regions undergoing neurogenesis. During the earlier timelapse (22.5-30hpf) a considerable morphological delay was observed in VPA-treated embryos, with the OT of VPA-treated embryos at 30hpf (Fig 3A IV’) resembling that of control embryos at 22.5hpf (Fig 3A I). However, despite these alterations in morphology, no appreciable neurogenesis was observed in the region of the OT until 30hpf in control embryos (Fig 3B IV), and no appreciable neurogenesis was observed at all in VPA-treated embryos during the earlier timelapse (Fig 3B IV’). During the later timelapse (30-43.5hpf) (Fig 3C,D) developmental delay is again evident in the overall OT morphology of VPA-treated embryos when compared to control. Images of 30hpf control embryos (Fig 3C I), where the transition from a columnar shape to the rounder cells characteristic of differentiating neurons has begun, resemble those of 43.5hpf VPA-treated embryos (Fig 3C IV’), indicating a delay in this transition. This delay becomes more evident when comparing *Tg(NeuroD:tRFP*^*w68*^) images (Fig 3D), which show a substantial decrease in neurogenesis in the region of the OT in VPA-treated embryos compared to control embryos at all imaged time points. Note: Discrepancies in *Tg(NeuroD:tRFP*^*w68*^) expression at 30hpf for the second timelapse (Fig 3D IV) compared to the same time point during the first timelapse (Fig 3B IV) are due to delays imposed from time away from a controlled incubation environment as the timelapse progressed.

**Figure 3.**
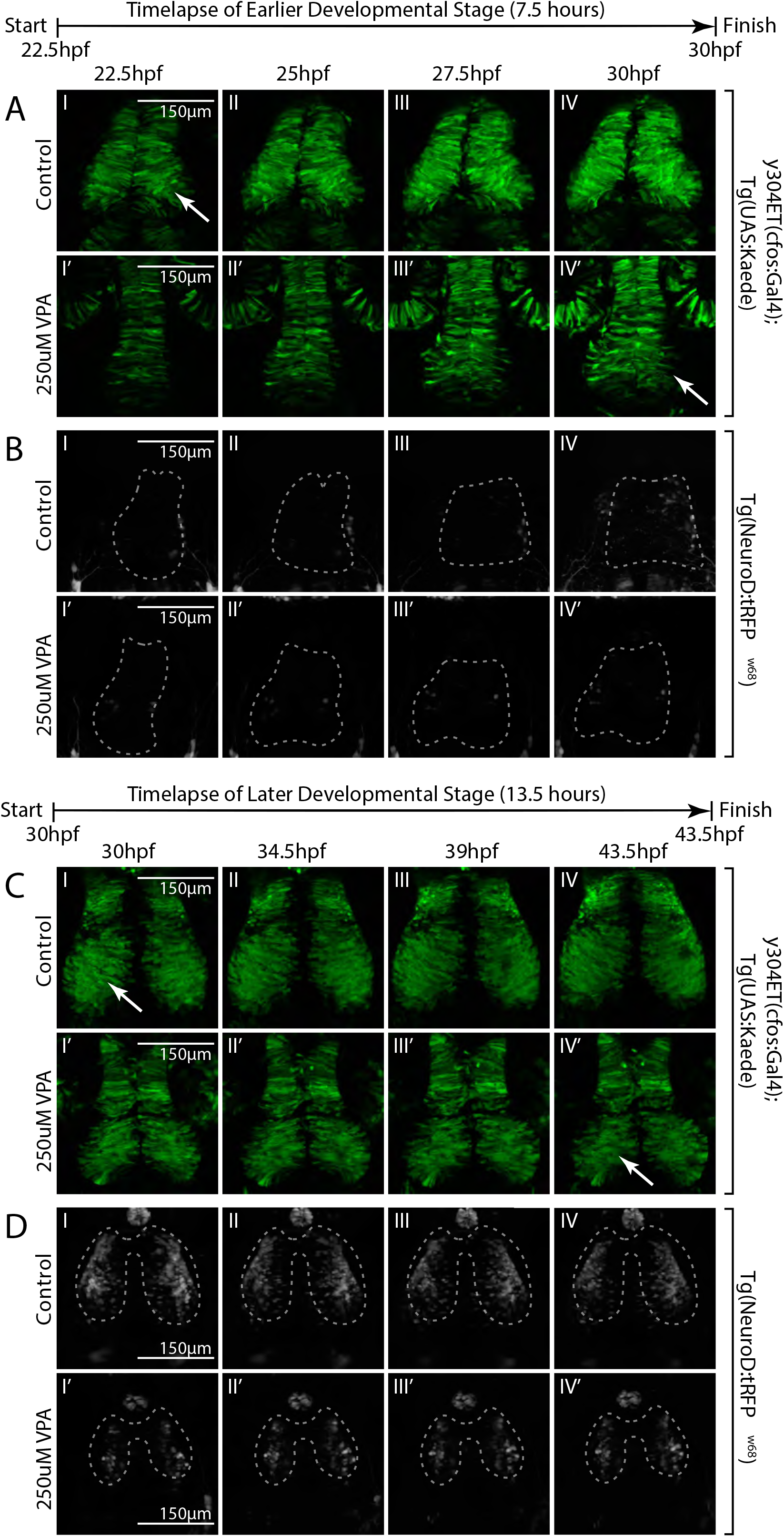
Timelapse imaging between 22.5-30hpf and between 30-43.5hpf shows a delay in neuronal differentiation and specification in VPA-treated embryos. (A) Timelapse imaging of VPA-treated and control *y304Et(cfos:Gal4); Tg(UAS:Kaede)* embryos at 22.5hpf-30hpf show that neuroepithelium proliferation (columnar cells) and early neuron generation/differentiation (shorter, rounder cells) is lagging in treated embryos (I’-IV’) and is more comparable (arrow, IV’) to controls at earlier time points (arrow, I). (B) At 22.5hpf-30hpf, no appreciable neuronal specification is seen in either control (I-IV) or treated embryos (I’-IV’) as determined by lack of fluorescence in the *Tg(NeuroD:tRFPw68)* line. (Dashed outlines=OT location). (C) At 30hpf-43.5hpf, neuroepithelium proliferation and neuron generation/differentiation continues to lag in treated embryos (arrow, IV’) when compared to controls (arrow, I). (D) During this time, more neurons become specified in controls (I-IV) when compared to VPA-treated embryos (I’’-IV”). Fluorescence in controls condenses into the shape of the OT as the timelapse progresses (I-IV). No increase in fluorescence is apparent for the treated embryos through the timelapse duration. (I’-IV’). Discrepancies in developmental stages between the movies of the two time periods of the same transgenic line result from delays imposed from time away from a controlled incubation environment as the timelapse progressed.

To visualize the long-term effects of VPA on OT neurogenesis we imaged *Tg(NeuroD:tRFP*^*w68*^) control and VPA-treated embryos daily from 24-120hpf (Fig 4) using the same treatment paradigm as previously discussed. Consistent with the timelapses (Fig 3), no appreciable neurogenesis occurred in the region of the OT in either control or VPA-treated embryos at 24hpf (Fig 4A I, B I). However, differences began to emerge at 48hpf with VPA-treated embryos (Fig 4B II) exhibiting weaker *NeuroD* expression than control embryos (Fig 4A II) in the OT. Interestingly, at 72hpf *NeuroD* expression appears more widespread in VPA-treated (Fig 4B III) than control larvae (Fig 4A III). This is consistent with the previously described scRNA-seq data (Suppl. Fig 2) suggesting that as the OT matures, *NeuroD* expression is restricted to only a subset of OT cell types, thus indicating that widespread *NeuroD* expression in the OT of VPA-treated larvae at 72hpf is likely indicative of a developmental delay. Restriction of *NeuroD* expression is eventually observed in VPA-treated larvae at both 96hpf and 120hpf; however, VPA-treated larvae lack the scattered neuropil expression and expression in arborization field 7 (AF7) (Fig 4A V arrow), a pretectal nucleus shown to be involved in hunting behavior ^40-42^, seen in control larvae (Fig 4A IV-V, IV’-V’). These results show that while delayed restriction of *NeuroD* expression indicates neural specification does eventually occur in VPA-treated larvae, it remains to be determined if they are specifying into the proper neuronal subtypes since some neuronal structures do not form in treated larvae.

**Figure 4.**
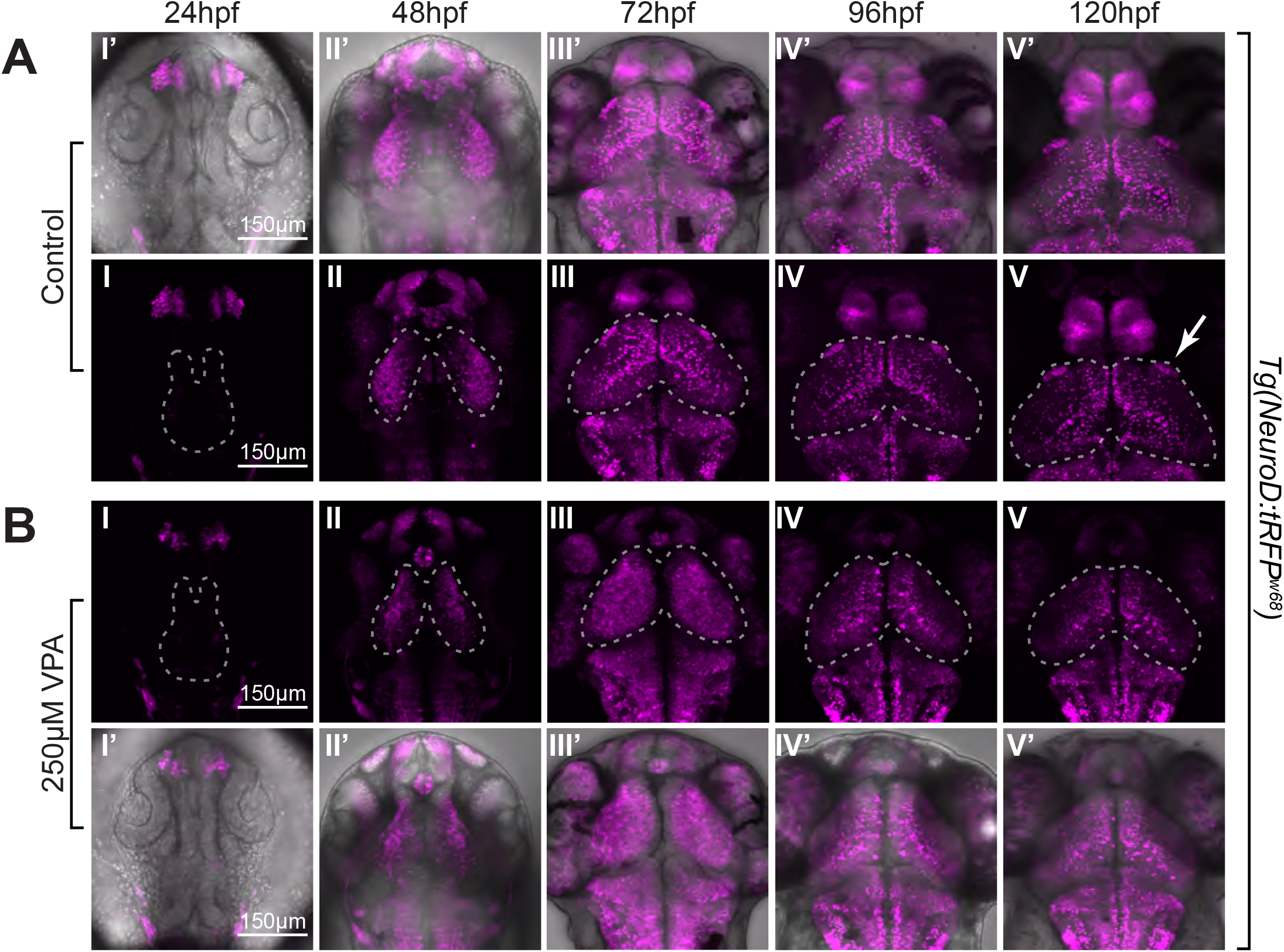
*Tg(NeuroD:tRFPw68)* imaging during the first 5 days of development reveals differences in neuronal maturation between control and treated embryos. Similar to Figure 2, we noticed no appreciable neuronal specification (as assessed by tRFP expression) in control (A: I, I’) or treated embryos (B: I, I’) at 24hpf, and more neuronal specification in control (A: II, II’) versus treated embryos (B: II, II’) at 48hpf. (Dashed outlines=OT location). As development progressed from 72-120 hpf, tRFP began to be restricted to subsets of neurons in control larvae (A: I-V; I’-V’). Instead, in treated larvae, tRFP expression was detected throughout the OT at 72hpf (B: III), which most likely resembles a time point between 48-72hpf in control embryos (not imaged). This expression becomes more restricted at 96-120hpf (B: IV,V,IV’,V’), suggesting that although neuronal specification lags or slows down in treated embryos, it does eventually occur. Whether the OT neurons in treated embryos are specified to become the same subclasses of neurons as in the control embryos remains to be determined. In addition, treated larvae lack expression in AF7 (arrow,V) and scattered neuropil expression seen in controls. Note: Panels I’-V’ in both A and B are overlays of fluorescent images on transmitted confocal projections.

### VPA Decreases Neurite Extension and Complexity in the Developing OT

To identify the role of VPA in preventing neuropil formation, we next investigated neurite extension and complexity within the OT of VPA-treated and control larvae. Since axonal tracts and synapses compose much of the neuropil, we utilized photoconversion of *Kaede* to visualize projections from a single neuron. Valproic acid-treated and control *y304Et(cfos:Gal4); Tg(UAS:Kaede)* embryos underwent the previously discussed treatment paradigm and were incubated in the dark to prevent photoconversion. Kaede was then photoconverted in individual neurons at 72, 96, and 120hpf to determine neurite length and branching. Valproic acid-treated larvae displayed neurons with shortened extensions into the neuropil and less complex branching at each time point measured (Fig 5 A-C I’-III’; Suppl. Videos 14-19). Lack of neurite extension into the neuropil, and decreased complexity, supports findings from the full-week *y304Et(cfos:Gal4); Tg(UAS:Kaede)* dosing experiments (Fig 2) where VPA-treated larvae exhibited little to no neuropil formation.

**Figure 5.**
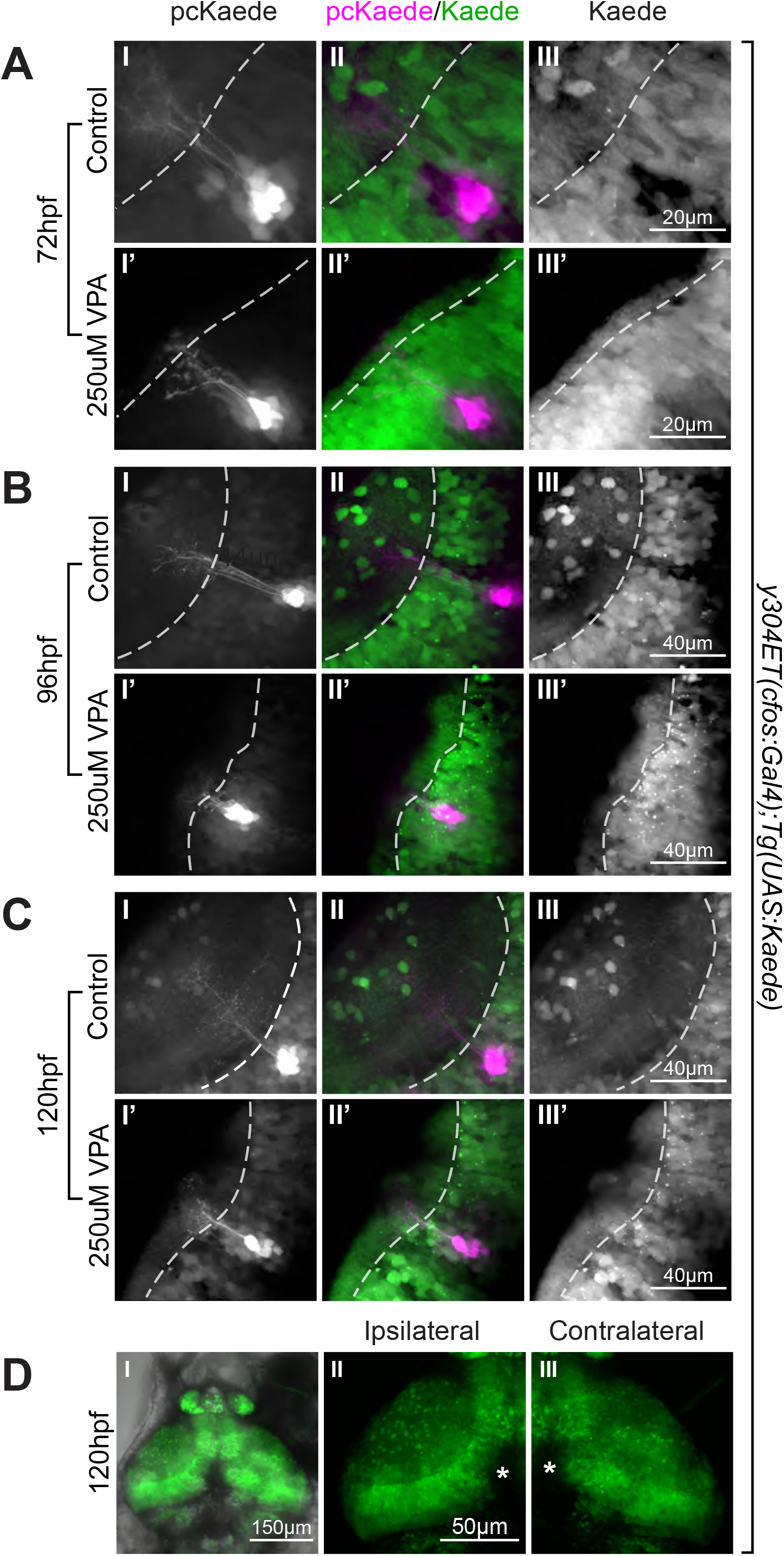
Photoconversions of neurons in the OT of VPA-treated embryos reveal shortened and less complex neuronal projections. (A-C) Photoconversion of Kaede from green to red (pseudo-colored magenta) was carried out in small groups of neurons in the *y304Et(cfos:Gal4); Tg(UAS:Kaede)* line to visualize projections from individual neurons. At 72hpf, 96hpf, and 120hpf, neurite growth is stunted in VPA treated larvae (I’-III’), compared to control larvae (I-III), resulting in shorter and less complex neurite projections; therefore, a smaller neuropil. (Dashed line= neuropil boundary). (D) Unilateral enucleations (left), show that both ipsilateral (II) and contralateral (III) tecti are identical to each other, indicating that the status of retinotectal axons is not integral to the initial development of OT neuron projections. This suggests that effects in OT axonogenesis and dendritogenesis are due to VPA effects on the OT and not on possible VPA effects on retinotectal projections from the retina, as seen in other studies (reference). (*) in II and III indicate pigment spots concealing the structures below.

The mechanisms underlying OT development are not fully understood, and likely rely on input from various sensory systems. Previous studies have shown that VPA exposure can lead to almost complete loss of retinotectal projections in a dose dependent manner ^43^. To determine if the loss of retinotectal extensions from the eye due to VPA exposure is responsible for the lack of neuropil seen in the OT of VPA-treated embryos, we imaged *y304Et(cfos:Gal4); Tg(UAS:Kaede)* embryos at 120hpf following unilateral enucleation at 32hpf (Fig 5D). Comparison of ipsilateral (Fig 5DII) and contralateral (Fig 5DIII) tectal lobes showed no apparent difference in neuropil formation. These results indicate that the shortened neurite extensions, decreased neurite complexity, and lack of neuropil observed in VPA-treated embryos are independent of VPA’s effect on retinotectal projections.

### VPA Effects Do Not Extend to All Neurons in the Embryo

After observing the effects of VPA on differentiation as well as on neurite extension and complexity within the OT, we next investigated if these effects were regionally localized to the OT or more generalized. To determine the effect of VPA in other regions, we imaged the spine of *Tg(NeuroD:tRFP*^*w68*^) VPA-treated and control embryos (Fig 6) utilizing the treatment paradigm discussed previously. Starting at 20hpf and continuing through 120hpf VPA-treated embryos showed a slight delay in development compared to their control counterparts, characterized by a decrease in complexity and extension of RoP, CaP and smn motor neurons, as well as almost complete loss of dorsal MiP motor neuron projections ^44^ even at later time points. However, while the effects of VPA were still observable in the spine, they were less extreme than the effects observed in the OT with VPA-treated embryos almost overcoming the delay by 120hpf (Fig 6 B IV-IV’). These results show that the effects of VPA treatment on the development of the nervous system extend beyond the OT, but that the OT is more susceptible than other neural structures.

**Figure 6.**
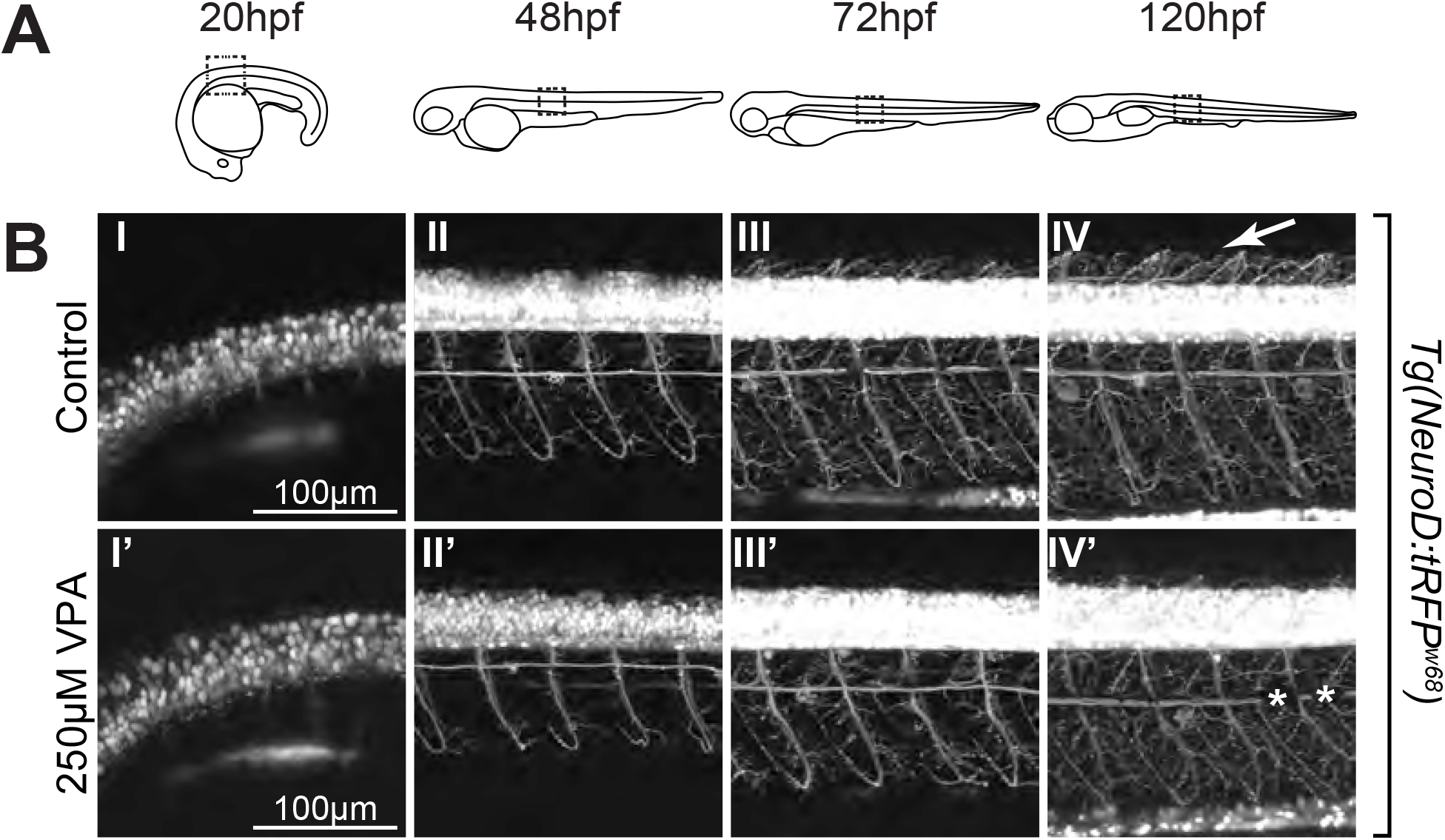
VPA effects do not extend to all neurons in the embryo. (A) Schematic of the imaging location at the different time points in embryos and larvae. Images of the spinal cord neurons in the *Tg(NeuroD:tRFPw68)* line indicate that neurogenesis (B) is occurring at the same time (∼20hpf) in control (I) and treated embryos (I’). Moreover, despite being somewhat delayed, axonogenesis and dendritogenesis in treated embryos (II’-IV’) does not look significantly different when compared to controls (II-IV) from 48-120hpf. The only major difference is the absence of dorsal projections of MiP motor neurons (arrow, IV) in the treated larvae. However, it is currently unknown if these projections are able to form at later time points. (*) in IV’ indicate pigment spots concealing the structures below. *Note: Due to high fluorescence level in the neuronal cell bodies, gamma has been adjusted (*.*60) uniformly on all images in panel B to allow for visualization of neuronal projections*.

### The Critical Window of OT Susceptibility to VPA-Treatment Includes Timepoints Prior to 72hpf

We next aimed to determine the critical window during which the OT is highly affected by VPA treatment. Since *NeuroD1* expression began to sequester to specific subpopulations at 72hpf (Fig 4A III), indicating completion of initial neurogenesis, and was fully sequestered by 96hpf (Fig 4A IV’), we identified timepoints prior to 54hpf or prior to 78hpf as possibilities for the OT critical window. To test this theory, embryos were dosed following the same paradigm used previously except that initial VPA treatment was delayed until 54hpf and 78hpf respectively. Embryos dosed starting at 54hpf (Fig 7B I’-V’) exhibited the phenotype previously observed in full week treatments (Fig 2) as quickly as 18 hours following exposure (Fig 7A III’), with almost complete loss of the neuropil at 120hpf (Fig 7A V’). When this experiment was repeated with treatment starting at 78hpf (Fig 7B I’-V’) the VPA-induced phenotype was greatly reduced. Although at 120hpf (Fig 7B V’) a delay was evident, larvae showed partial neuropil formation, the presence of some superficial interneurons (SINs), and segregation of the periventricular layer (PVL) from the neuropil, all of which was not seen in embryos exposed to full week treatments.

**Figure 7.**
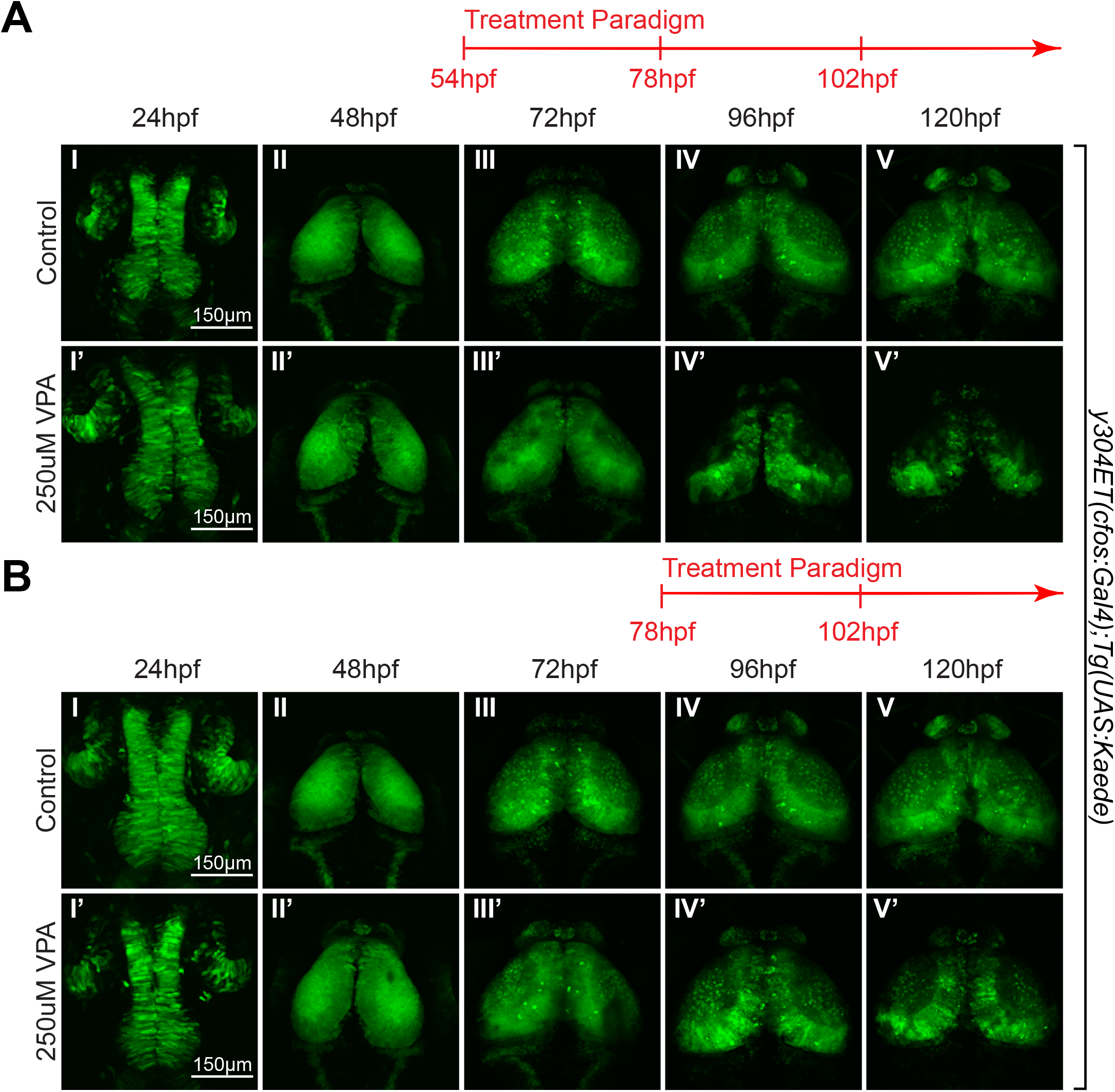
The critical window of OT susceptibility to VPA-treatment includes timepoints prior to 72hpf. (A) Treatment of *y304Et(cfos:Gal4); Tg(UAS:Kaede)* embryos starting at 54-120hpf, still resulted in neuropil development defects in the treated embryos (I’-V’) compared to controls (I-V). Therefore, the critical window of susceptibility must be after 54hpf (B) A subsequent trial in which VPA treatment was initiated at 78hpf resulted in a significantly reduced phenotype in treated larvae (IV’-V’). Therefore, the critical window of susceptibility is determined to be timepoints prior to 72hpf.

To further validate the existence of a critical window, and to determine at what time point the embryos could recuperate from the effects of VPA, we imaged *y304Et(cfos:Gal4); Tg(UAS:Kaede)* embryos before and after single-day VPA treatments starting at 6hpf, 30hpf, and 54hpf. Embryos exposed to single-day VPA treatment at all tested timepoints exhibited developmental delays with embryos treated starting at 6hpf (Fig 8B) and 30hpf (Fig 8C) showing delays comparable to embryos treated for a full-week (Fig 2). Treatment from 54-78hpf caused delays evident as soon as 96hpf (Fig 8D IV) and which continued through 120hpf (Fig 8D V), although these delays were much less severe than embryos treated at earlier timepoints. These results suggest that the critical window for the OT’s sensitivity to VPA includes timepoints prior to 78hpf, with timepoints prior to 54hpf exhibiting heightened sensitivity to VPA treatment where single day doses are sufficient to induce phenotypes comparable to week-long treatment.

**Figure 8.**
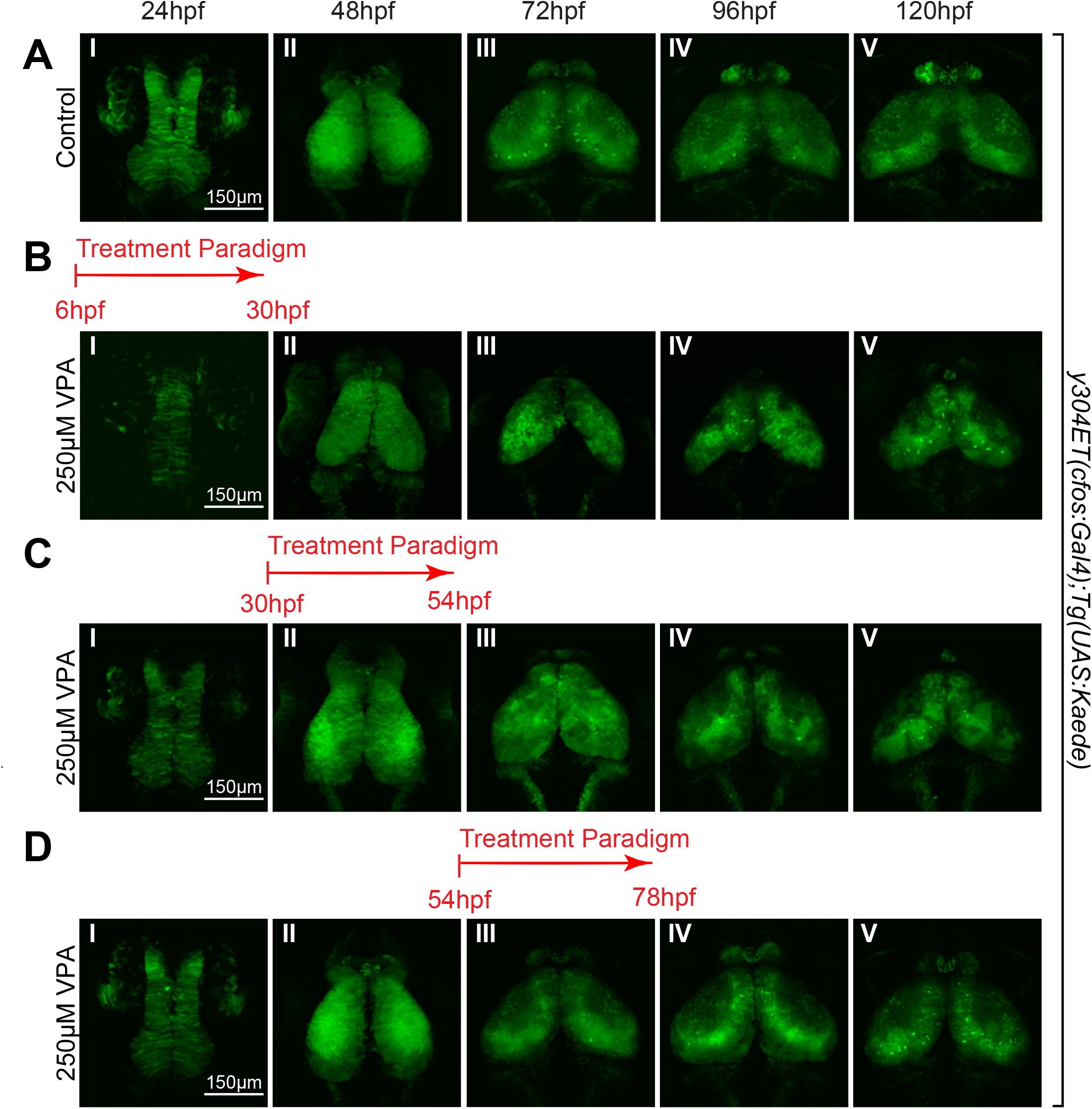
Single 24hr VPA treatment prior to 78hpf still result in developmentally delayed OTs. Short, 24hr VPA treatments were administered during three different time-windows followed by replacement of the solution with EM. Daily imaging of control (A) and 250µM VPA-treated *y304Et(cfos:Gal4); Tg(UAS:Kaede)* embryos show that one 24hr VPA-treatment at 6hpf-30hpf (B) or 30hpf-54hpf (C) still leads to a smaller OT and neuropil (B:II-V; C:III-V) and that the development, despite the shorter treatment time, did not recuperate. (D) When the VPA-treatment was applied between 54-78hpf, OT development was less affected than in previous treatments (B,C); however the OT was still underdeveloped and exhibited a smaller neuropil compared to control embryos (A).

### Upregulation of the Nrf2 Antioxidant Response System and Increased GSH_Tot_ Fails to Protect Against the Effects of VPA on OT Development

Previous research done in mouse embryonic teratoma cells identified oxidative stress as a possible mechanism for the negative effects of VPA exposure and demonstrated that upregulation of the nuclear factor-erythroid factor 2-related factor 2 (nrf2) antioxidant response system can ameliorate the negative effects of VPA on neurogenesis *in vitro* ^45^. To investigate the interplay between the nrf2 antioxidant response and VPA in the OT, we administered the nrf2-inducer 3H-1, 2-dithiol-3-thione (D3T) prior to VPA treatment to upregulate expression of specific redox-related genes throughout the embryo. The genes of interest included: *hmox1a, hmo1b, gclc, nqo1, and gstp1*. To verify that D3T was inducing increased expression of these genes, embryos were exposed to D3T for 12 hours (6hpf-18hpf). Following 12 hours of D3T treatment, embryos were collected and prepared for RT-qPCR analysis. When compared to control embryos, *gclc* and *gstp1* showed significantly increased expression (Fig 9A). Both *gclc* and *gstp1* play a direct role in reduced glutathione (GSH) synthesis and usage, an important molecule in the antioxidant response. Other genes trended toward an increase but were not significantly different.

**Figure 9.**
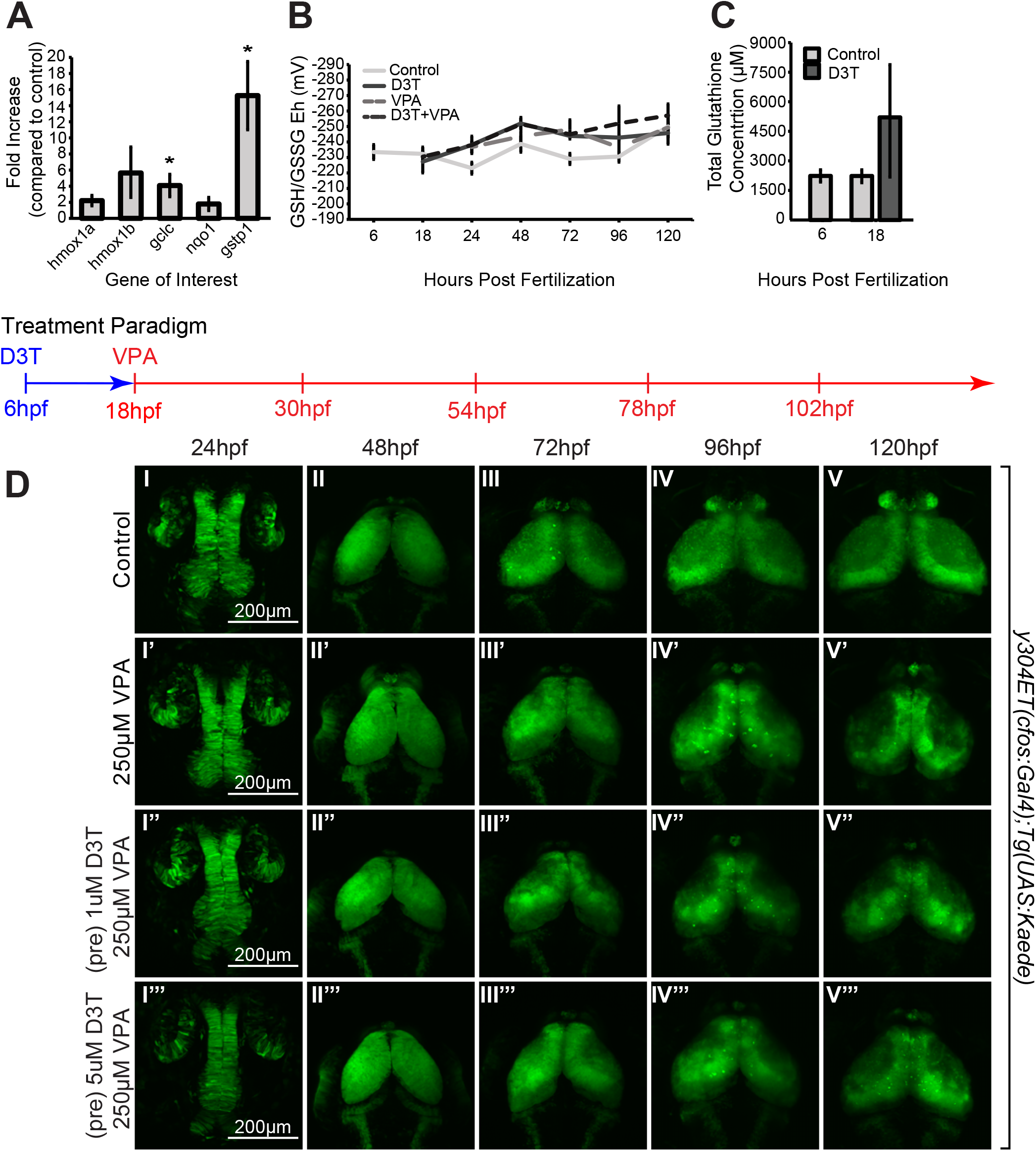
Induction of genes downstream of the antioxidant-response transcription factor, nrf2, do not protect against VPA effects in the OT. (A) Treatment of embryos from 6-18hpf with 5µM D3T induces expression of some genes (*Gclc* and *Gstp1* that play a direct role in reduced glutathione (GSH) synthesis and usage) downstream of *nrf2* transcription factor gene in total embryos. (B) However, pretreatment for 12 hours with 5µM D3T before incubation with 250µM VPA did not significantly change the redox potential when compared to VPA only treated embryos at any of the tested time points, even though D3T-treated embryos showed increased total glutathione (reduced and oxidized) concentration at 18hpf compared to control embryos (C). A reducing trend is seen in all treated embryos compared to controls (B); however, these trends are not significant. Data points (A-C) represent n=3 pools of 30 embryos. Mean ±SEM is plotted. (*) denotes p≤0.05 compared to control. (D) Pretreatment for 12 hours with 1μM (I’’-V’’) or 5μM (I’’’-V’’’) D3T before incubation with 250μM VPA provides no protection in the OT compared to embryos treated with VPA alone (I’-V’) despite upregulation of *nrf2* target genes.

To measure redox states after VPA exposure, embryos were treated with 250µM VPA beginning at 18hpf, and media was replaced every 24 hours until 5dpf. Pools of 30 embryos were collected at 24hpf and each following day up to 5dpf to be derivatized for HPLC analysis. Expecting an oxidizing shift following VPA treatment compared to control embryos, we also treated embryos with the nrf2 inducer, D3T at 6hpf, prior to VPA treatment. At 18hpf, D3T was removed and 250µM VPA was added to some of the embryos following the dosing and collection scheme mentioned above. Interestingly, there was no significant difference in the measured redox potentials between control embryos and embryos treated with D3T, VPA, or D3T+VPA from 6hpf-5dpf, although a slight trend toward a reducing redox potential was observed (Fig 9B). Although D3T pretreatment did increase total GSH concentration, there was no significant change in the redox state when compared to VPA treatment alone (Figs 9B and 8C). To determine the effect of D3T pretreatment on OT phenotype following VPA exposure, *y304Et(cfos:Gal4); Tg(UAS:Kaede)* control, VPA, 1µM D3T+VPA, and 5µM D3T+VPA embryos were imaged every 24 hours from 24-120hpf (Fig 9D) following the dosing paradigm just discussed. At 120hpf, no improvement was seen between embryos treated with only VPA (Fig 9D V’) and those pre-treated with either 1µM (Fig 9D V’’) or 5µM D3T (Fig 9D V’’’). Interestingly, pre-treatment with D3T appeared to amplify the negative effects of VPA at 96hpf and 120hpf with embryos exhibiting a smaller neuropil than VPA-treated embryos (Fig 9D IV’-V’, IV’’, V”, IV’’’, V’’’). Additionally, timelapse imaging during the initial OT critical period also revealed no rescue in D3T pretreated embryos (Suppl. Fig 5; Suppl. Videos 11-13). These results suggest that D3T pretreatments are not sufficient to ameliorate VPA-induced delays in OT development.

## CONCLUSIONS

Timelapse movies of *y304Et(cfos:Gal4); Tg(UAS:Kaede)* embryos showed that starting around 22hpf the neuroepithelium, from which the OT will be derived, undergoes extensive proliferation resulting in neurons that begin to be specified soon after, around 28-30hpf *(Tg(NeuroD:tRFPw68) movies)*. Embryos treated with VPA are still able to undergo proliferation; however, the marker for specification (tRFP driven by the *neuroD* promoter) seems to be delayed. From our current experiments, it is difficult to tell if the rate of proliferation or the actual neuronal specification is delayed, but it would be interesting in the future to follow up this observation with timelapse-imaging of embryos that have a nuclear reporter or have been given a BrdU pulse. Previous studies have shown varying results as to whether VPA inhibits or promotes proliferation ^46-49^, suggesting that dose and treatment length may alter outcomes. One study ^50^ of zebrafish embryos treated with VPA identified upregulation of several miRNAs, including those predicted to regulate cell cycle genes. Among those cell cycle regulating genes, *cdkn1a, tp53* are broadly expressed throughout the different neuronal subclasses at 7dpf in our scRNA-seq data ^39^, while *wee1, cdk2* and *chek1* are specifically expressed in one developing neuronal population ^39^ (Suppl. Fig 2B). It will be worthwhile in the future to determine how these genes regulate OT proliferation during development.

As mentioned above, we found that in control embryos neuronal specification of the OT begins around 28-30hpf *(Tg(NeuroD:tRFPw68) movies)*. At 72hpf the tRFP reporter expression is expanded and then restricted at days 4 and 5 to a subset of OT populations. Instead, tRFP expression onset in VPA treated embryos is delayed from 28-30hpf to 72hpf, and then progresses, although slower than controls, in days 4 and 5. This delay in neuronal specification is consistent with findings from neuronal cultures treated with VPA ^45^. It is important to keep in mind that although the delay in neuronal specification is resolved, it is currently unclear if all the neuronal subclasses are subsequently generated, and if the number for each subclass remains the same. Jacob and colleagues ^51^ showed that VPA treatment of zebrafish embryos resulted in downregulation of the proneural gene *ascl1b* (associated with Notch signaling) via the blockade of HDAC1 inhibitor that is required for its expression. Could these same genes play a role in OT neurogenesis? Our scRNA-seq data shows that *HDAC1* and *HDAC4* are indeed present in 7dpf OT neurons. Although *ascl1b* is not highly expressed in any of the OT neuronal populations at this stage, *ascl1a* and *Her* genes upstream of *ascl1* in the Notch pathway are present in the developing populations (Suppl. Fig 2C-D). This suggests that HDAC1, HDAC4 and the Notch pathway might play a role in OT neuronal specification.

Even though neurogenesis occurs, although greatly delayed, in VPA treated embryos, we found that neurite extension and branching is severely affected at 72-120hpf. This phenotype does not seem to be dependent on the potential lack of retinotectal projections from the retinal ganglion cells, which were previously shown to be affected by VPA^43^. Early removal of one of the eyes does not appear to significantly affect the contralateral tectal neuropil at 5dpf, indicating that the neurite phenotype we see is most likely due to VPAs effect on OT neurons. Therefore, a more plausible explanation for this phenotype is that VPA affects either axon guidance molecules or cytoskeletal proteins during neuronal development. Interestingly, Aluru et al, 2013^50^, found that VPA treatment of zebrafish embryos also led to the downregulation of several siRNAs whose targets are axon guidance genes, which is consistent with our observations. Although VPA treatment affects OT neuron development, the development of the majority of the spinal cord neurons (with exception of MiP motor neurons whose dorsal projection appear to not extend during the observed time period)^44^ are generally slightly delayed but able to catch up. This suggests that the development of OT and spinal cord neurons is overseen by genes that have non-overlapping functions.

Finally, we wanted to determine if the effects of VPA on OT development were due to oxidative stress. The transcription factor nrf2 plays a key role in the expression of many detoxifying and antioxidant enzymes as well as GSH biosynthesis enzymes to resist oxidative stress during differentiation^52,53^. Under normal conditions, nrf2 goes through a controlled degradation process dependent on Kelch like ECH-associated protein 1 (Keap1), a nrf2-specific adaptor protein for the Cul3 ubiquitin ligase complex ^54-56^. Past studies have shown that various chemicals, including D3T, are nrf2 activators through their interactions with KEAP1 and cause the dissociation of KEAP1/nrf2 complex to allow nrf2 to subsequently accumulate in the nucleus and regulate gene expression ^45,55,57,58^. This response system was hypothesized to be a protective mechanism against VPA exposure during zebrafish OT development. However, in the current study, we show that D3T does not have a protective effect in the OT. Although the GSHTot was increased and various *nrf2*-regulated genes showed a significant increase in expression after D3T pre-treatment, this heightened concentration of GSH and GSSG did not prevent malformation in the OT following VPA exposure. The RT-qPCR and HPLC data represent whole embryos and not solely the OT, so it is unclear what direct effect D3T pretreatments may have on the OT. The results may be different if the redox potential and gene expression exclusively of the OT were measured, which are likely the focus of future experimentation. It would be expected from the data shown in this study that both the redox potential and gene expression remain the same when compared to VPA-treated embryos.

Alternatively, it could be that the expression of nrf2 itself is not high enough early in development when we exposed embryos to VPA. One study showed no significant upregulation of nrf2-regulated genes (namely *gstp1*) nor nrf2 activation at 8-10hpf after exposure to a wide range of chemicals without the addition of exogenous nrf2 ^58^. The exposure to an oxidant this early in development may come prior to any antioxidant response pathway being robust enough to ensure a functional, beneficial, response and may cause malformation, disruption of protein function, and ROS-mediated damage with limited protection available. Since VPA-induced effects could not be resolved by pretreatment with D3T, there is a strong possibility that the observed phenotypes are not due to oxidative stress, but rather to a possible inhibitory effect on HDAC and an aberrant effect on gene expression profiles. As previously mentioned HDAC1 and HDAC4 are present in OT at 7dpf and their specific status in the OT following VPA treatment should be followed up in the future.

In conclusion, our study utilized VPA to perturb development of the OT for the purpose of understanding the mechanisms underlying proper development of the OT. Our results indicate that VPA treatment induces gross morphological alterations as well as negatively impacts neurogenesis, axonogenesis, and dendritogenesis. Additionally, our findings identify for the first time a critical period for OT patterning, neurogenesis, and neurite extension during which the OT is especially vulnerable to VPA treatment. Finally, this work provides a foundation for research into mechanisms driving OT development as well as the relationship between the OT, VPA, and ASD.

## ACKNOWLEDGEMENTS

We would like to thank Ella Bleak for help with microscopy work; Tanya Finken, Larissa Larson, and Liv Olson for excellent care of the fish facility.

## FIGURE LEGENDS

**Supplementary Figure 1. OT imaging reveals the ability of pH to modulate the effects of VPA in the OT**. Daily images of control (I-V) and treated (I’-V’) *y304Et(cfos:Gal4); Tg(UAS:Kaede)* embryos with 250µM VPA dissolved in embryo media pH6.6 (A), pH7.2 (B), and pH7.8 (C). Treated embryos were continuously exposed to VPA solution from 6-120hpf. (A) VPA-treated embryos incubated at pH6.6 (I’-IV’) exhibited strong phenotypes such as dysmorphic, twisted tecti as soon as 48hpf (arrow, II’) and no neuropil formation at 96hpf. Moreover, they did not survive for imaging at 120hpf. (B) VPA-treated embryos incubated at pH7.2 (I’-V’) did not exhibit twisting of the tecti as seen at pH6.8 (A), but they displayed a smaller OT and lacked neuropil formation at 72-120hpf. (C) VPA-treated embryos incubated at pH7.8 (I’-V’) presented a slightly weaker phenotype, characterized by an increase in OT size (V’), compared to embryos incubated at pH7.2 (B) or pH6.8 (A). *Note: OT development in control embryos was unaffected by pH*.

**Supplementary Figure 2. Presence of genes of interest in the OT at 7dpf determined by scRNA-seq data**. Relative expression of various genes of interest within cellular subtypes of the OT. Point size indicates percentage of cells expressing the gene within a population. Point color indicates average gene expression within a population. (A) At 7dpf all *NeuroD* genes were expressed in the OT with *NeuroD1* found in the greatest number of OT clusters. Note that although *NeuroD1* can be found in many clusters, it is largely sequestered to OT7 and OT19 by 7dpf. (B) Expression of various cell cycle genes reveal that *cdkn1a* and *tp5d* are found throughout the OT at 7dpf. In contrast, *wee1, cdk2, and chek1*, are largely found in OT10, a cluster identified as developing neuronal ^39^. (C) Expression profiles show that both *HDAC1* and *HDAC4* are present in the OT at 7dpf. *Ascl1a*, an important downstream target of the Notch pathway, is also found in the OT at 7dpf, although *ascl1b* does not appear to be present. (D) Various *her* genes, additional downstream targets of the Notch pathway, show widespread expression between clusters and a high percentage of cells expressing within those clusters at 7dpf.

**Supplementary Figure 3. GSH (reduced glutathione), GSSG (oxidized glutathione), and GSHTot (reduced and oxidized) during development**. GSH (A), GSSG (B), and GSHTot (C) concentrations over the course of embryonic development (6-120hpf) for control, 5µM D3T, 250µM VPA, and 5µM D3T + 250µM VPA-treated embryos. D3T+VPA-treated embryos received a 12-hour D3T pretreatment from 6-18hpf, following which D3T+VPA and VPA-treated embryos were continuously exposed to VPA from 18-120hpf. Concentration was determined by protein quantification assay and HPLC. By 5dpf all treated groups showed increased GSH (A) and GSHTot (C) compared to controls. All data points represent n=3 pools of 30 embryos each. Data plotted as mean ±SEM.

**Supplementary Figure 4. Extended dosing paradigm demonstrates that prolonged exposure to high concentrations of D3T alters OT development**. Daily images of control (I-V), DMSO (I’-V’), and 10µM D3T-treated (I’’-V’’) *y304Et(cfos:Gal4); Tg(UAS:Kaede)* embryos. Treated embryos were continuously exposed to DMSO (I’-V’) or 10μM D3T (I’’-V’’) from 6-120hpf. At 48hpf embryos treated with 10μM D3T (II’’) showed increased columnar neuroepithelial cells (arrow, II’’), and an overall decrease in size, when compared to both control (II) and DMSO embryos (II’). This delay became increasingly more apparent with D3T-treated larvae showing severe malformation in the OT at 120hpf (V’’). DMSO-treated embryos (I’-V’) did not exhibit noticeable deviations from control embryos (I-V) at any measured timepoints.

**Supplementary Figure 5. Timelapse immediately after D3T pretreatment indicates that D3T does not ameliorate the effects of VPA on OT development during the critical period**. Images captured at regular intervals from a 20-42.5hpf timelapse of *y304Et(cfos:Gal4); Tg(UAS:Kaede)* control (I-IV), 250μM VPA (I’-IV’) and 5μM D3T+250µM VPA-treated (I’’-IV’’) embryos. D3T+VPA-treated embryos received a 12-hour D3T pretreatment from 6-18hpf following which D3T+VPA and VPA embryos were continuously exposed to VPA from 18-42.5hpf. Pretreatment with 5μM D3T (I’’-IV’’) failed to remedy the effects of VPA on OT development when compared with control (I-IV), and VPA-treated (I’-IV’) embryos. At 20hpf D3T+VPA embryos (arrow, I’’) lack proliferation of the posterior neuroepithelium seen in control embryos (arrow, I). By 42.5hpf D3T+VPA embryos exhibit persistent columnar neuroepithelial cells (arrow, IV’’) in contrast to control embryos which display more rounded cells indicative of differentiation (arrow, IV).

**Supplementary Video 1: *y304Et(cfos:Gal4); Tg(UAS:Kaede)* Timelapse from 22.5hpf-30hpf, Control**

**Supplementary Video 2: *y304Et(cfos:Gal4); Tg(UAS:Kaede)* Timelapse from 22.5hpf-30hpf, 250µM VPA**

**Supplementary Video 3: *y304Et(cfos:Gal4); Tg(UAS:Kaede)* Timelapse from 30hpf-43.5hpf, Control**

**Supplementary Video 4: *y304Et(cfos:Gal4); Tg(UAS:Kaede)* Timelapse from 30hpf-43.5hpf, 250µM VPA**

**Supplementary Video 5: *Tg(NeuroD:tRFPw68)* Timelapse from 22.5hpf-30hpf, Control**

**Supplementary Video 6: *Tg(NeuroD:tRFPw68)* Timelapse from 22.5hpf-30hpf, 250µM VPA**

**Supplementary Video 7: *Tg(NeuroD:tRFPw68)* Timelapse from 30hpf-43.5hpf, Control Supplementary Video 8: *Tg(NeuroD:tRFPw68)* Timelapse from 30hpf-43.5hpf, 250µM VPA**

**Supplementary Video 9: *Tg(NeuroD:tRFPw68)* Timelapse from 30hpf-43.5hpf, Control - angle #2**

**Supplementary Video 10: *Tg(NeuroD:tRFPw68)* Timelapse from 30hpf-43.5hpf, 250µM VPA-angle #2**

**Supplementary Video 11: *y304Et(cfos:Gal4); Tg(UAS:Kaede)* Timelapse from 20hpf-42.5hpf: 250µM VPA**

**Supplementary Video 12: *y304Et(cfos:Gal4); Tg(UAS:Kaede)* Timelapse from 20hpf-42.5hpf: 5µM D3T**

**Supplementary Video 13: *y304Et(cfos:Gal4); Tg(UAS:Kaede)* Timelapse from 20hpf-42.5hpf: Control**

**Supplementary Video 14: 3D Rotations of photoconverted *y304Et(cfos:Gal4); Tg(UAS:Kaede)* neurons. 250µM VPA 3dpf**

**Supplementary Video 15: 3D Rotations of photoconverted *y304Et(cfos:Gal4); Tg(UAS:Kaede)* neurons. 250µM VPA 4dpf**

**Supplementary Video 16: 3D Rotations of photoconverted *y304Et(cfos:Gal4); Tg(UAS:Kaede)* neurons. 250µM VPA 5dpf**

**Supplementary Video 17: 3D Rotations of photoconverted *y304Et(cfos:Gal4); Tg(UAS:Kaede)* neurons. Control 3dpf**

**Supplementary Video 18: 3D Rotations of photoconverted *y304Et(cfos:Gal4); Tg(UAS:Kaede)* neurons. Control 4dpf**

**Supplementary Video 19: 3D Rotations of photoconverted *y304Et(cfos:Gal4); Tg(UAS:Kaede)* neurons. Control 5dpf**

## Notes

**Funding** This work was supported by the NICHD: R15HD095737 and internal university funds.

**Conflict of interest disclosure** The authors declare that the research was conducted in the absence of any commercial or financial relationships that could be construed as a potential conflict of interest.

### Competing Interest Statement

The authors have declared no competing interest.

