## Supplementary figures and images for "Valproic Acid Affects Neuronal Specification and Differentiation During Early Optic Tectum Development of Zebrafish"

### Supplemental Figure1

Supplementary Figure 1

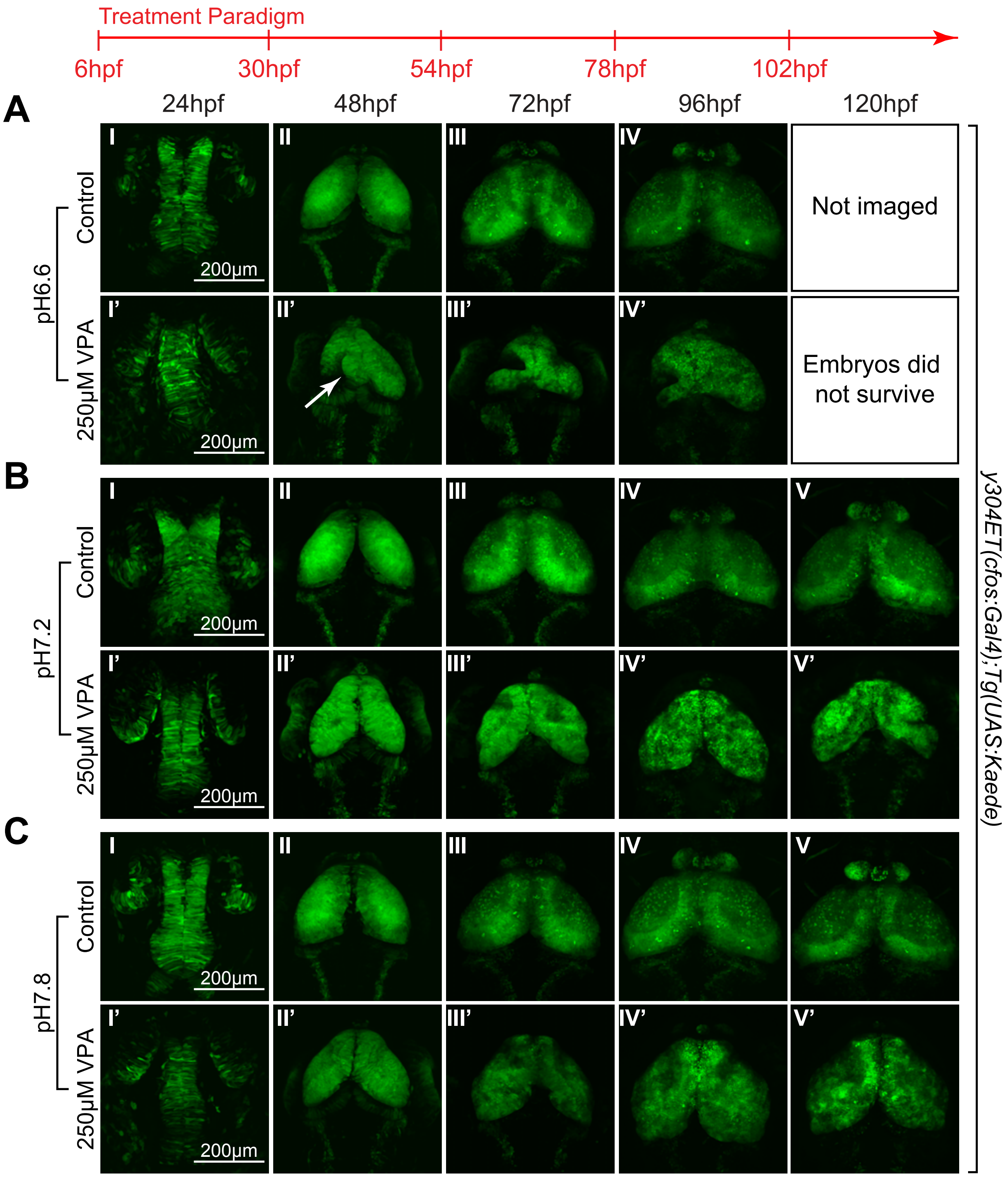

### Supplemental Figure2

Supplementary Figure 2

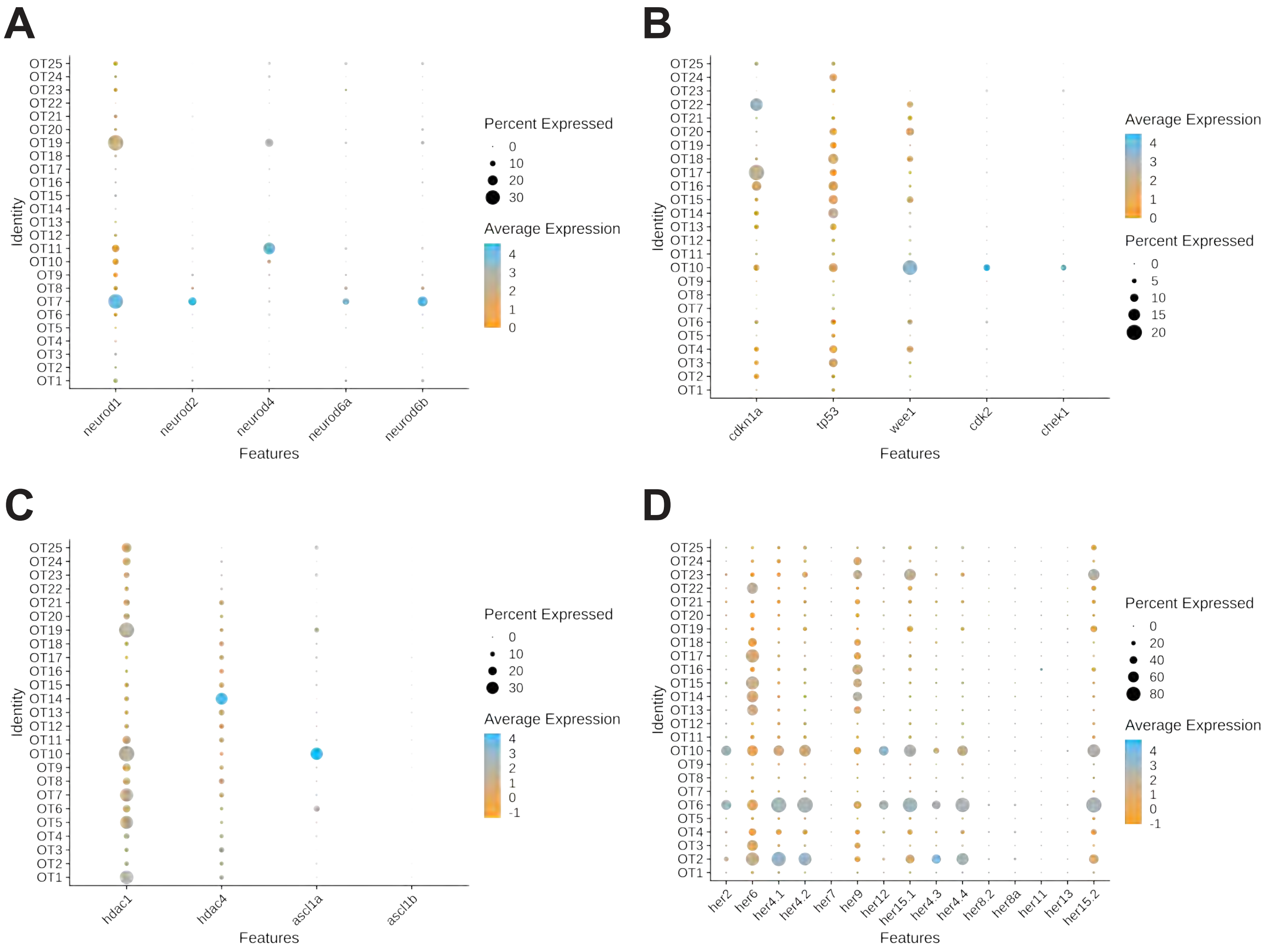

### Supplemental Figure3

Supplementary Figure 3

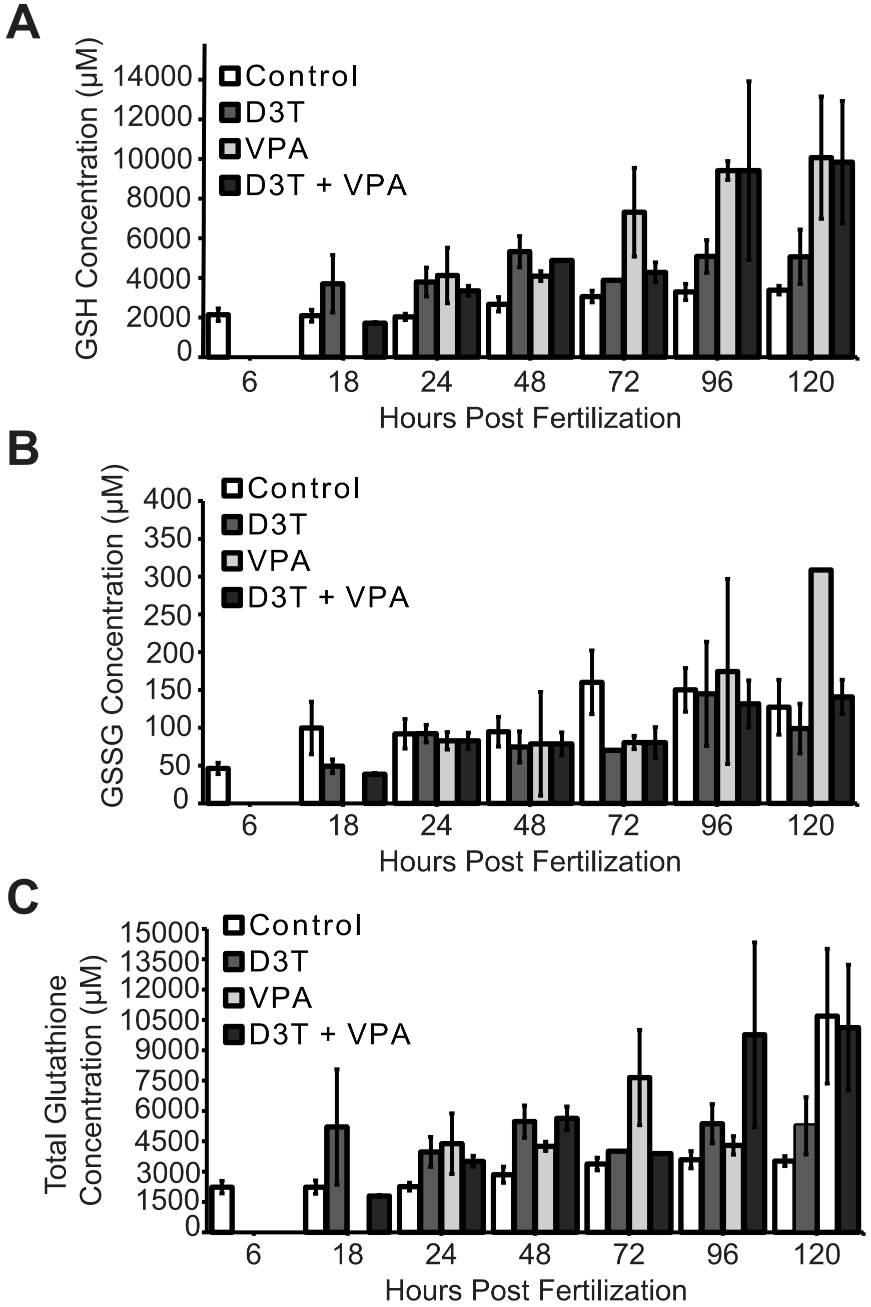

### Supplemental Figure4

Supplementary Figure 4

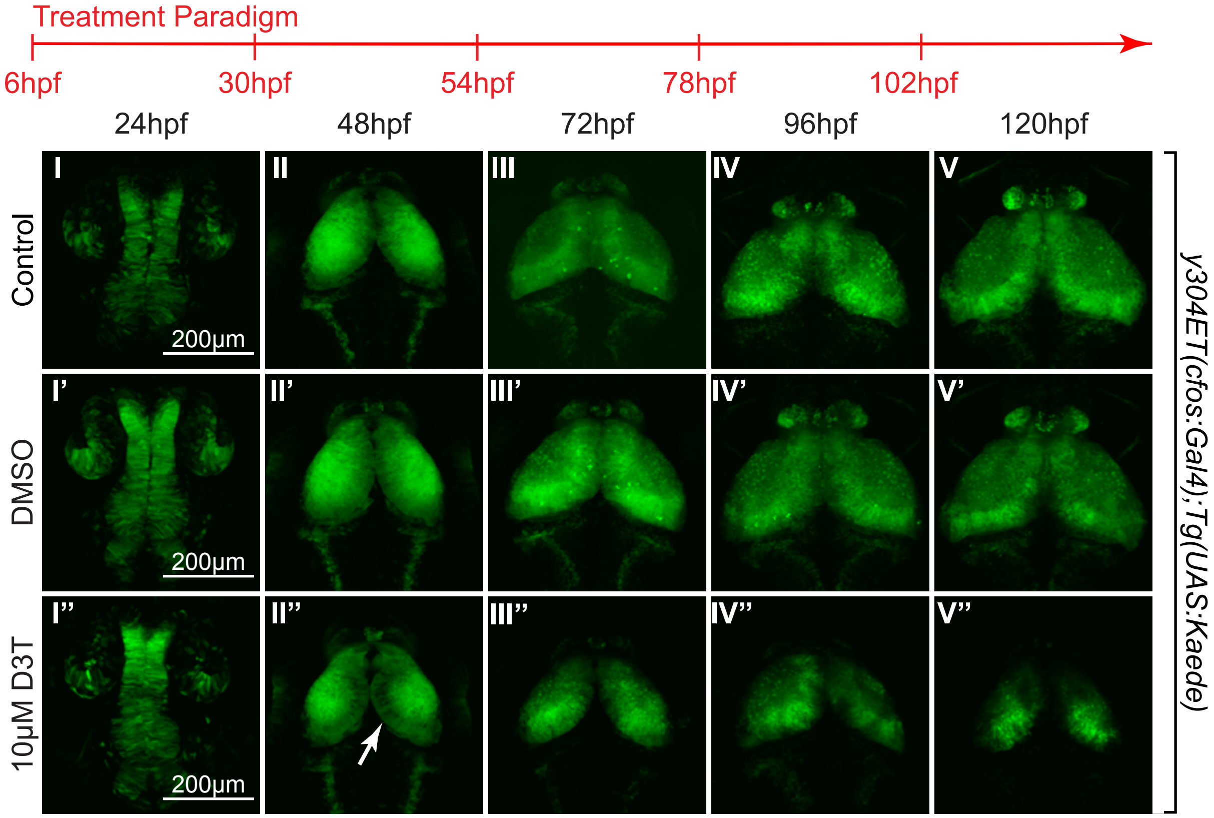

### Supplemental Figure5

Supplementary Figure 5

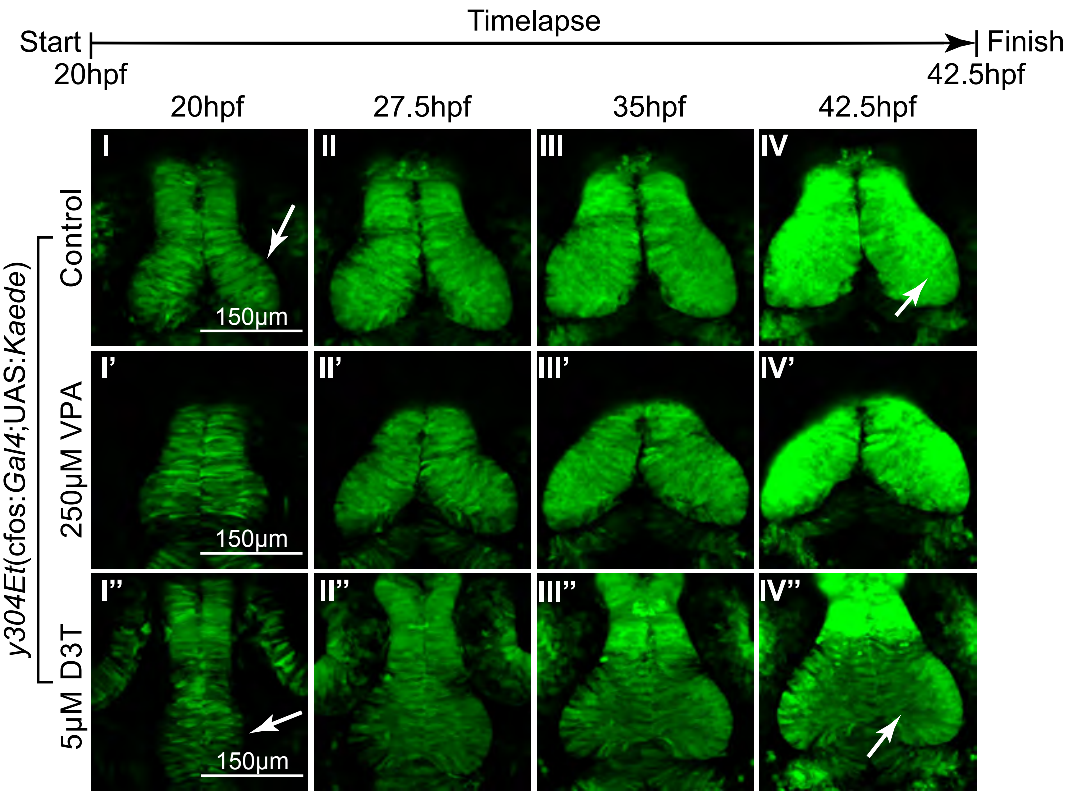
